# Inactivation of GalU leads the cell wall-associated polysaccharide defect to reduce the susceptibility to bacteriolytic agents in *Enterococcus faecalis*

**DOI:** 10.1101/2020.11.20.391417

**Authors:** Jun Kurushima, Haruyoshi Tomita

**Author notes:** Corresponding author: Haruyoshi Tomita.

## Abstract

Enterococcal plasmid-encoded bacteriolysin Bac41 is a selective antimicrobial system that is considered to provide a competitive advantage to *Enterococcus faecalis* cells that carry the Bac41-coding plasmid. The Bac41 effector consists of the secreted proteins BacL1 and BacA, which attack the cell wall of the target *E. faecalis* cell to induce bacteriolysis. Here, we demonstrated that *galU*, which encodes UTP-glucose-1-phosphate uridylyltransferase, is involved in susceptibility to the Bac41 system in *E. faecalis*. Spontaneous mutants that developed resistance to the antimicrobial effects of BacL1 and BacA were revealed to carry a truncation deletion of the C-terminal 288–298 a.a. region of the translated GalU protein. This truncation resulted in the depletion of UDP-glucose, leading to a failure to utilize galactose and produce the enterococcal polysaccharide antigen (EPA), which is expressed abundantly on the cell surface of *E. faecalis*. This cell surface composition defect that resulted from *galU* or EPA-specific genes caused an abnormal cell morphology, with impaired polarity during cell division and alterations of the limited localization of BacL1. Interestingly, these mutants conferred reduced susceptibility to beta-lactams, despite their increased susceptibility to other bacteriostatic antimicrobial agents and chemical detergents. These data suggest that a complex mechanism of action underlies lytic killing, as exogenous bacteriolysis induced by lytic bacteriocins or beta-lactams requires an intact cell physiology in *E. faecalis*.

**Importance:** Cell wall-associated polysaccharides of bacteria are involved in various physiological characteristics. Recent studies demonstrated that the cell wall-associated polysaccharide of *Enterococcus faecalis* is required for susceptibility to bactericidal antibiotic agents. Here, we demonstrated that a *galU* mutation resulted in resistance to the enterococcal lytic bacteriocin Bac41. The *galU* homologue is reported to be essential for biosynthesis of species-specific cell wall-associated polysaccharides in other Firmicutes. In *E. faecalis*, the *galU* mutant lost the *E. faecalis*-specific cell wall-associated polysaccharide EPA (enterococcal polysaccharide antigen). The mutant also displayed reduced susceptibility to antibacterial agents and an abnormal cell morphology. We firstly demonstrated that *galU* was essential for EPA biosynthesis in *E. faecalis*, and EPA production might underlie susceptibility to lytic bacteriocin and antibiotic agents by undefined mechanism.

## Introduction

*Enterococcus faecalis* is a Gram-positive opportunistic pathogen that is a causative agent of several infectious diseases, such as urinary infections, bacteremia, endocarditis, and others (1). This organism belongs to the lactic acid bacteria and is also a microbial resource that produces various bacteriocins (2). Bacteriocins are antimicrobial proteins or peptides that are produced by bacteria and are considered to play a role in bacterial competition within the microbial ecological environment (3). Gram-positive bacterial bacteriocins are classified according to their structures or synthetic pathways (4). Class I bacteriocins, which are referred to as lantibiotics, are heat-stable peptides that contain non-proteinogenic amino acids modified by post-translational modifications(5). Class II bacteriocins are also heat-stable peptides, but they are synthesized without any post-translational modification (6). In *E. faecalis*, beta-hemolysin/bacteriocin (cytolysin), which is the major virulence factor, and enterocin W belong to the class I bacteriocins (7–9). On the other hand, most enterococcal bacteriocins, including Bac21, Bac31, Bac32, Bac43, Bac51 and others, belong to the class II peptides(6, 8, 10–15). Class III bacteriocins are proteinaceous heat-liable bacteriocins that differ from the class I or II peptides. Therefore, bacteriocins of this class are often referred to as bacteriolysins. Two enterococcal bacteriolysins, enterolysin A and Bac41, have been identified to date (16, 17).

The bacteriolysin Bac41 was originally found in the conjugative plasmid pYI14 of *E. faecalis* clinical strain YI14 and is widely distributed among clinical isolates including vancomycin resistant enterococci (VRE) (17–19). The activity of this bacteriocin is specific towards *E. faecalis* and not other *Enterococcus* or bacterial species (20). The genetic element of Bac41 consists of *bacL*_*1*_, *bacL*_*2*_, *bacA*, and *bacI*. BacL_1_ and BacA are the effectors responsible for the antimicrobial activity of this bacteriocin against *E. faecalis*. BacL2 is involved in the expression of antimicrobial activity as transcriptional positive regulator (17, 21). BacI is required for the self-protection of Bac41-producing *E. faecalis* cells from the antimicrobial activity of BacL_1_ and BacA. Both the BacL_1_ and BacA molecules contain conserved peptidoglycan hydrolase domains, suggesting that these molecules attack the target’s cell wall (17). Endopeptidase activity against the *E. faecalis* cell wall has been detected experimentally in BacL_1_ (22). However, the peptidoglycan degrading activity of BacA has not been detected, and the enzymatic function of this molecule remains unknown. BacL_1_ binds specifically to the nascent cell wall or to cell division loci via its C-terminal SH3 repeat domain (20). The resulting cell growth inhibition protects against the binding of BacL_1_ and markedly reduces susceptibility to bacterial killing by Bac41 effectors. It is important to note that the BacL_1_ cell wall degradation process is essential but not sufficient to kill the bacteria; bacteriolysis requires the additional action of the undefined BacA.

In this paper, we isolated a spontaneous mutant of *E. faecalis* that exhibited resistance to the toxicity of BacL_1_ and BacA to further understand the bacteriolytic mechanism of the Bac41 system. Genomic variant analysis showed that an intact *galU* gene was essential for susceptibility to BacL_1_ and BacA. Furthermore, we demonstrated that *galU* is also essential for galactose fermentation and for the synthesis of the enterococcal polysaccharide antigen, which is required for maintaining the normal cell morphology and for susceptibility to antimicrobial agents.

## Results

### Inactivation of *galU* results in decreased susceptibility to Bac41

To identify the target *E. faecalis* cell factor(s) involved in Bac41-mediated bacterial killing, spontaneous mutants that are resistant to the Bac41 effectors BacL_1_ and BacA were obtained. We carried out a soft-agar bacteriocin experiment in which the susceptible bacterial strain *E. faecalis* OG1S was inoculated into and grown in THB agar (0.75 %), and a mixture of recombinant BacL_1_ and BacA proteins (25 ng of each protein) was spotted onto the *E. faecalis*-containing THB agar. After incubation at 37 °C overnight, a clear growth inhibition zone was formed in the area where the recombinant proteins had been spotted (22). Notably, additional incubation resulted in the occurrence of small colonies within the growth inhibitory zone. These colonies were considered to be spontaneously resistant mutants, so we isolated these colonies as candidates that may carry a genetic mutation related to bacterial killing by Bac41. To identify the genetic mutation that caused this spontaneous Bac41 resistance, we performed whole-genome sequencing and a variant analysis of the mutant strains. The contigs from the genome sequence of the parent strain *E. faecalis* OG1S and the isogenic mutant were obtained by next generation sequencing and were mapped onto the reference strain *E. faecalis* OG1RF (accession No. NC_017316). One of these spontaneous mutants, sr #4, carried the point mutation C862T within the *galU* (annotated as *cap4C* in the OG1RF genomic information) coding sequence (Fig. 1A). The wild-type *galU* gene consists of 897-bp, and its translated product GalU is 298 a.a. in length. The mutation found in sr #4 was a nonsense mutation and resulted in the truncation of the GalU protein by a C-terminal deletion of 288–298 a.a. (Fig. 1B). Complementation via the introduction of the intact *galU* gene carried by the expression vector p*galU* restored the mutant strain’s susceptibility to Bac41 activity to the level of the parent strain (Fig. 1C). In addition, a liquid phase bacteriolytic assay also demonstrated that the spontaneous mutant (referred to as *galU*^−^ hereafter) could still grow in the presence of recombinant BacL_1_ (rBacL_1_) and BacA (rBacA), while the wild-type strain underwent considerable lysis (Fig. 1D). The complementation of the *galU* mutation via the expression of intact *galU in trans* (*galU*^−^/p*galU*) but not the control vector (*galU*^−^/Vec) restored the strain’s susceptibility to bacteriolysis to the same degree as the wild-type strain. The viability of the *galU*^−^ and *galU*^−^/Vec strains exhibited a >100-fold increase in comparison to the WT and *galU*^−^/p*galU* strains after treatment with rBacL_1_ and rBacA (5 μg/ml each) for 4 h (Fig. 1E). Therefore, these results strongly demonstrated that the truncated mutation of the *galU* gene resulted in resistance to the antimicrobial activity of Bac41.

**Figure 1.**
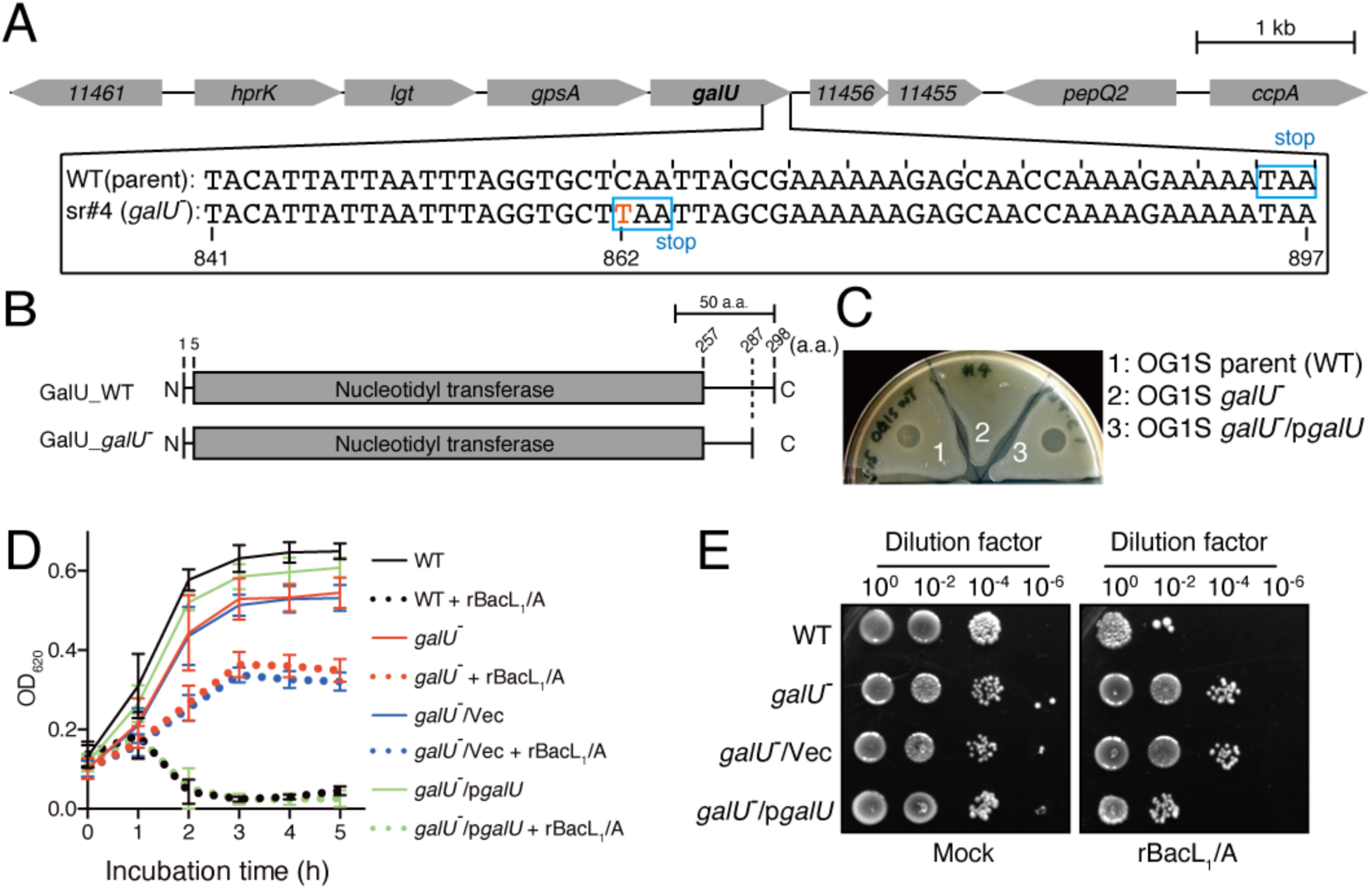
Genetic mutation found in the strain spontaneously resistant to the antimicrobial activity of Bac41. **(A)** Nucleotide sequence alignment of the *galU* (OG1RF_11457; Gene ID: 12287400) genes of the *E. faecalis* OG1S parent strain (upper) and the OG1S isogenic mutant strain (sr#4) that spontaneously acquired resistance to the antimicrobial activity of Bac41 (lower). Scale bar, 1 kb. (B) Amino acid sequence alignment of the translation products of the *galU* genes of the *E. faecalis* OG1S parent strain (upper) and the OG1S isogenic mutant strain (lower). Scale bar, 50 a.a.. (C) A mixture of recombinant BacL_1_ -;His and BacA-His proteins (25 ng each) was spotted onto THB soft agar (0. 75 %) containing the indicator strain *E. faecatis* OG1S (Parent, 1), the spontaneously resistant strain with the *galU* mutation (*galU*^−^, 2), and the *galU*^−^ strain complemented via the trans-expression of wild-type GalU from the *pgalU* plasmid *(galU*^−^*pgalU*, 3). The plate was incubated at 37 °C for 24 h, and the formation of halos was evaluated. (D) Overnight cultures of *E. faecatis* strains, including OG1S (WT), the *galU* mutant *(galU-)*, the vector control strain of *galU-(galU-Nec)* and the complemented *gall.i* strain *(gall.ilpgalU)*, were inoculated into fresh THB broth at a 5-fold dilution. A mixture of recombinant BacL,-His (5 μg/ml) and BacA-His (5 μg/ml) was added to the bacterial suspension and incubated at 37 °c. The turbidity was monitored during the incubation period. The data for each case are presented as the mean ±S.D. (error bars) of three independent experiments. (E) The *E. faecalis* strains were treated with rBacL_1_ and BacA (5 μg/ml each) at 37 °c for 4 h, as described in panel D. The bacterial suspensions were serially diluted 100-fold with fresh THB and then spotted onto a THB agar plate, followed by incubation overnight. Colony formation was evaluated as a measure of bacterial viability.

### The enterococcal *galU* is essential for the fermentation of galactose and the production of cell wall-associated polysaccharides

The *galU* gene encodes UDP-glucose phosphorylase (UDPG::PP, EC: 2.7.7.9), which generates UDP-glucose from a UTP molecule and glucose-1-phosphate (23). The homologues of this gene are universally conserved not only in prokaryotes but also in eukaryotes (24). It has been reported extensively that among both Gram-positive and Gram-negative bacteria, this enzymatic activity of GalU is essential for the carbohydrate metabolism that is involved in multiple bacterial physiological pathways; however, the precise role of GalU in *E. faecalis* remains undefined. UDP-glucose, which is the product of GalU, is utilized for galactose fermentation via the Leloir pathway in lactic acid bacteria (Fig. 2A) (25). We tested whether the strain that possesses an inactivated *galU* has the ability to undergo galactose fermentation (Fig. 2B). The wild-type strain and the complemented strain had the ability to ferment both glucose and galactose. In contrast, the *galU* mutant strain and the *galU*^−^/Vec strain could utilize glucose but not galactose. In addition, the in-frame deletion mutants of *galU* that were constructed via site-directed mutagenesis also failed to utilize galactose in the two different lineages OG1RF and FA2-2 (Supplementary figure S1). These data clearly indicated that GalU produces UDP-glucose in *E. faecalis* and in other species and also suggested that the C-terminal truncation of GalU in the *galU*^−^ strain resulted in the complete loss of its UDP-glucose phosphorylase (UDPG::PP) activity.

**Figure 2.**
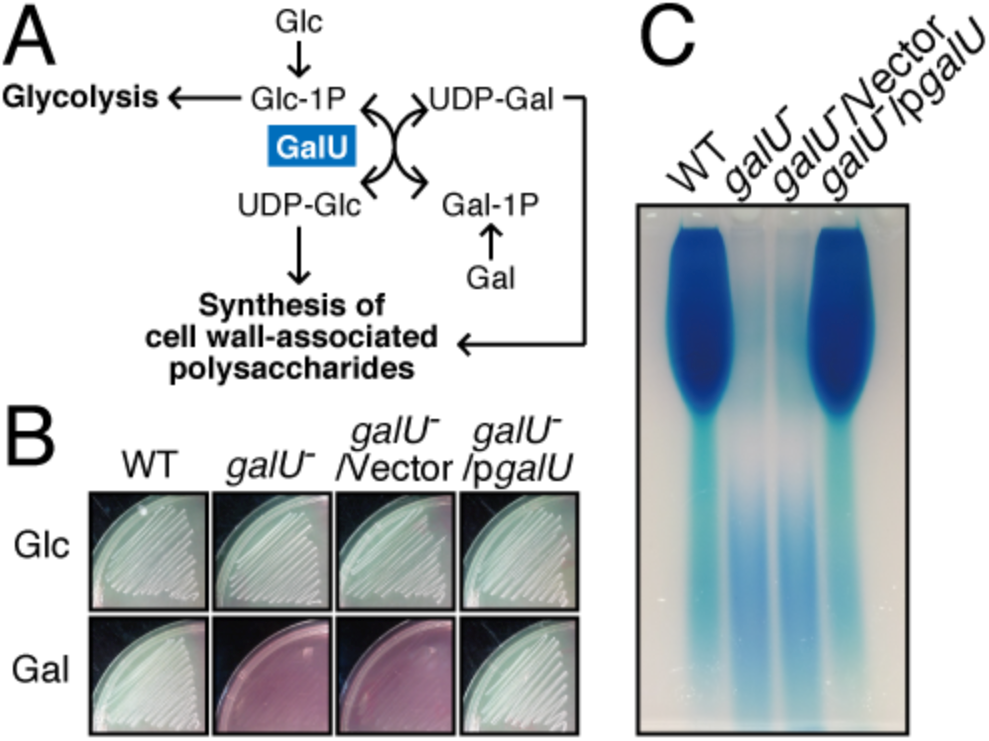
*ga/U* is essential for galactose fermentation and cell surface polysaccharide production. (A) The Leloir pathway of galactose metabolism is illustrated. GalU generates UDP-glucose (UDP-Glc) from glucose-1-phosphate (Glc-1P) and ATP. (B) The E *faecalis* wild-type, *galU*^−^, *galu*^−^/*Vec* and *galU*^−^*lpga/U* strains were grown in HI agar medium supplemented with phenol red as a pH indicator and glucose (Glc) or galactose (Gal) as the fermentation source. (C) The *E. faecalis* wild-type, *galU*^−^, *galU*^−^/*Vec* and *galU*^−^*lpga/U* strains were grown in THB broth supplemented with glucose, and the cell wall-associated polysaccharides were prepared as described in the Materials and Methods section. The resulting polysaccharides were separated via 10 % acrylamide gel electrophoresis, followed by staining with the Stains-All reagent.

In addition to its role in galactose fermentation, GalU and its product, UDP-glucose, are essential for the biosynthesis of bacterial glycopolymers, such as LPS, teichoic acid and capsular polysaccharide, which are important structural components that determine the properties of the bacterial cell surface (26). For example, in *S. pneumoniae*, which has more than 90 capsular variants, GalU is essential for the biosynthesis of various types of capsular polysaccharide (Mollerach et al., 1998). The cell surface polysaccharide profiles of *E. faecalis* strains were previously demonstrated (27, 28). To investigate the effect of the *galU* mutation on capsular polysaccharide synthesis in enterococci, cell wall-associated polysaccharides were prepared from the *E. faecalis* WT, *galU*^−^, *galU*^−^/Vec and *galU*^−^/p*galU* strains and subjected to polyacrylamide gel electrophoresis analysis, followed by visualization with cationic staining (Fig. 2C). The most abundant band, which corresponds to the enterococcal polysaccharide antigen (EPA), was observed in the wild-type OG1S and *galU*^−^/p*galU* strains. In contrast, the EPA had completely disappeared in the *galU*^−^ and *galU*^−^/vector strains. This result strongly suggested that the *galU* gene is essential for EPA production in *E. faecalis*.

Previous studies identified the specific EPA biosynthesis gene cluster *epaA–R* (27, 29, 30). Several *epa* gene mutants with altered EPA composition, such as mutants of *epaA, B, E, M, N, I, X* or *O*, displayed a pleiotropic phenotype that included abnormal cell morphology, defective biofilm formation, and increased susceptibility to antimicrobial agents and phages, as well as other phenotypes (30, 31). We investigated the effect of *galU* inactivation on cell morphology. *E. faecalis* is an ovococcal bacterium with an ellipsoid-cell shape, and its size is approximately 2 μm by 1 μm. Observation under a scanning electron microscope revealed that the *galU*^−^ strain showed an abnormal cell morphology with a rounded cell shape and a slightly larger size (~2 μm) (Fig. 3A). Fluorescent imaging using DAPI, which is a fluorescence probe for DNA, revealed that abnormal multinucleated cells were present in the *galU*^−^ population (Fig. 3B). The complemented strain *galU*^−^/p*galU* but not the control vector *galU*^−^/Vec showed the same morphological phenotype as the wild-type strain. These observations suggested that the truncation of the *galU* gene leads to a marked morphological defect.

**Figure 3.**
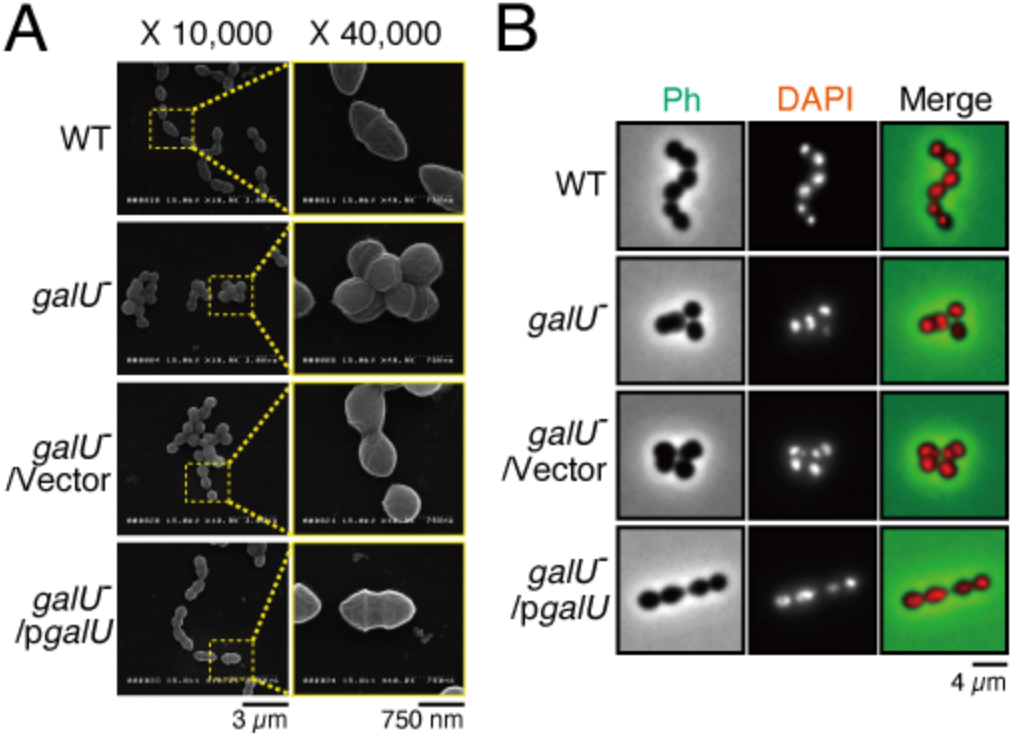
*ga/U* is essential for maintaining cell morphology. (A) The *E. faecalis* wild-type, *ga/U*^−^, *ga/U*^−^/*Vec* and *ga/U-lpga/U* strains were grown on cover glasses, followed by chemical fixation. The samples were subjected to osmium coating and analyzed under the scanning electron microscope. Scale bars, 3 *μm* (left) and 750 nm (right). (B) Overnight cultures of the *E. faecalis* wild-type, *ga/U*^−^, *ga/U*^−^/*Vec* and *ga/U-lpga/U* strains were diluted 5-fold with fresh THB broth and incubated, chemically fixed and mounted with DAPI for DNA visualization (red), followed by analysis via fluorescence microscopy. Phase contrast (Ph) is pseudocolored (green) in the merged image. Scale bar, 4 *μm*.

### The loss of cell wall-associated EPA does not block targeting and cell wall degrading of BacL_1_ to the cell wall

We previously reported that the peptidoglycan degradation activity of BacL_1_ is not sufficient but still essential for Bac41-mediated bacterial killing (22). We examined whether the observed resistance to Bac41 activity in the *galU*^−^ mutant results from decreased susceptibility of the cell wall to BacL_1_. Cell wall fractions prepared from the wild-type OG1S strain and the *galU*^−^, *galU*^−^/Vec or *galU*^−^/p*galU* strains were treated with recombinant BacL_1_, followed by incubation at 37 °C (Fig. 4A). The cell walls from the parent strain and the complemented strain were degraded to 52.8 % and 51.9 % of the initial amount after 6 hours of incubation, respectively. In contrast, the *galU*^−^- and *galU*^−^/Vec-derived cell walls were also degraded to 70.6 % and 68.5 %, respectively, by BacL_1_, although degradation rates were slightly attenuated compared to those of the parent strain and the complemented strain. In addition, the cell walls of *galU*^−^ and *galU*^−^/Vec also displayed reduced susceptibility to mutanolysin, which is a peptidoglycan hydrolase but has no capacity to induce definite bacteriolysis (Supplementary figure S2) (22). These observations suggested the possibility that cell wall of *galU* mutant is still susceptible to peptidoglycan hydrolase activity of BacL_1_. The binding of BacL_1_ via its C-terminal SH3 repeat domain, which is located in 329–380 a.a., is limited to the cell division loci of the target cell wall and is essential for the peptidase activity of BacL_1_ (22). To investigate whether the binding affinity of BacL_1_ for the *galU*^−^ cell wall was involved in attenuated susceptibility to degradation by BacL_1_, fluorescence-labeled recombinant BacL_1_ was incubated with *E. faecalis* cells and its localization was analyzed by fluorescent microscopy. In the parent strain and the complemented strain, BacL_1_ binding was limited to division loci, as reported previously (Fig. 4B, Supplementary figure S3A) (20). In *galU*^−^ mutant, BacL_1_ strongly bound to over the cells surface even more diffusely and was less limited to specific target loci, showing that the affinity of BacL_1_ for the cell wall the *galU*^−^ strain appeared to be even more increased compared to those of wild type. These observations suggested that attenuated susceptibility to BacL_1_ degradation is not likely cause of the development of Bac41 resistance in the *galU* mutant, even though the overall cell surface affinity for BacL_1_ was altered in the *galU^−^ mutant*.

**Figure 4.**
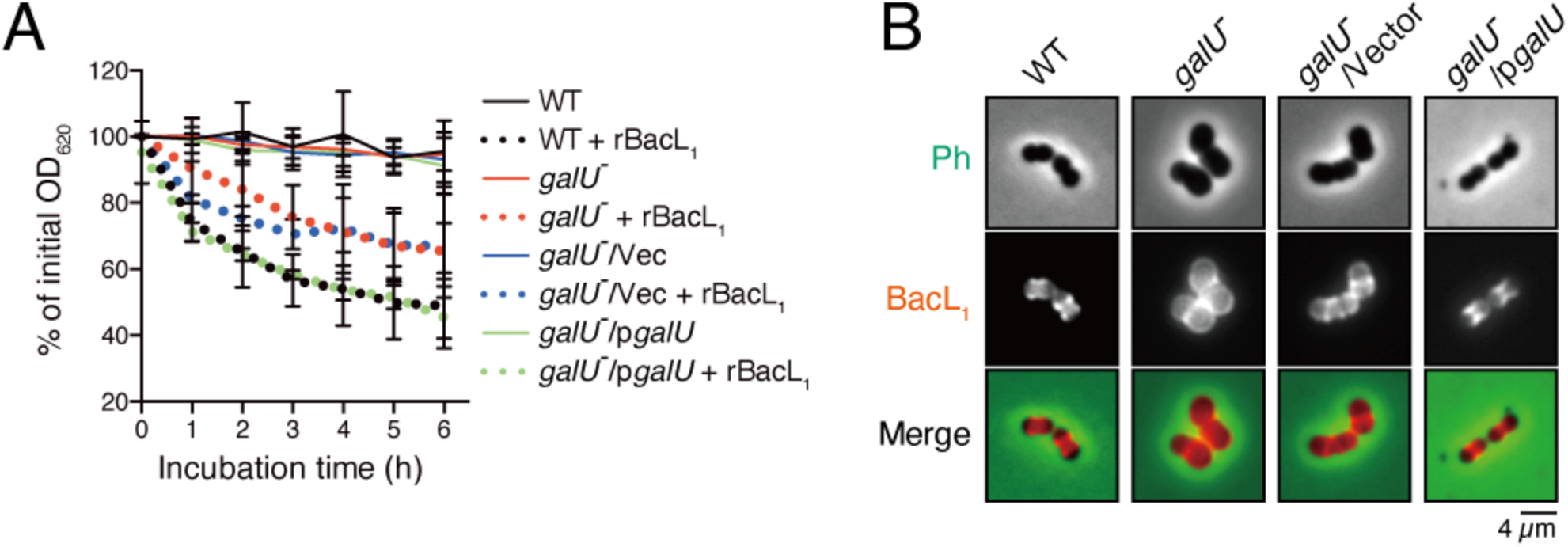
*ga/U* inactivation alters the interaction of BacL_1_ with the cell wall of *E. faecalis*. (A) Cell wall fractions prepared from *E. faecalis* wild-type (black), *ga/U*^−^ (red), *ga/U*^−^/*Vec* (blue) and *ga/U*^−^*lpga/U* (green) in exponential phase were diluted with PBS. Recombinant BacL_1_ (rBacL_1_, 5 μg/ml, dotted lines) was added to the cell wall suspension, and the mixture was incubated at 37 °C. The turbidity at 600 nm was quantified at the indicated times during incubation. The presented values are the percentages of the initial turbidity for the respective samples. The PBS-treated sample (mock) is presented as a negative control (straight lines). The data are presented as the mean ±S.D. (error bars) of four independent experiments. (B) Overnight cultures of the *E. faecalis* wild-type, *ga/U*^−^, *ga/U*^−^/*Vec* and *ga/U*^−^*lpga/U* strains were diluted 5-fold with fresh **THB** broth and incubated with Hilyte Fluor 555 fluorescent dye-labeled (red) BacL_1_ (5 μg/ml), followed by analysis via fluorescence microscopy. Phase contrast (Ph) is pseudocolored (green) in the merged images. Scale bar, 4 *μm*.

To investigate whether the phenotypes of the *galU*^−^ mutant resulted specifically from the loss of cell wall-associated polysaccharides, we constructed a deletion mutant of *epaB*, which is involved in the major EPA biosynthesis pathway (Fig. 5A) (31). The inactivation of *epaB* is suggested to result in an altered EPA structure and composition and reduced cell envelope integrity. The *epaB* mutant (*ΔepaB*) no longer produces EPA (30, 32), meanwhile it displayed normal galactose fermentation as like wild type (Supplementary figure S4). Notably, the *ΔepaB* clearly represented resistance to Bac41 as *galU*^−^ mutant (Fig 5B and C), suggesting that Bac41 resistance in *galU*^−^ mutant is mostly due to inability of EPA production.

**Figure 5.**
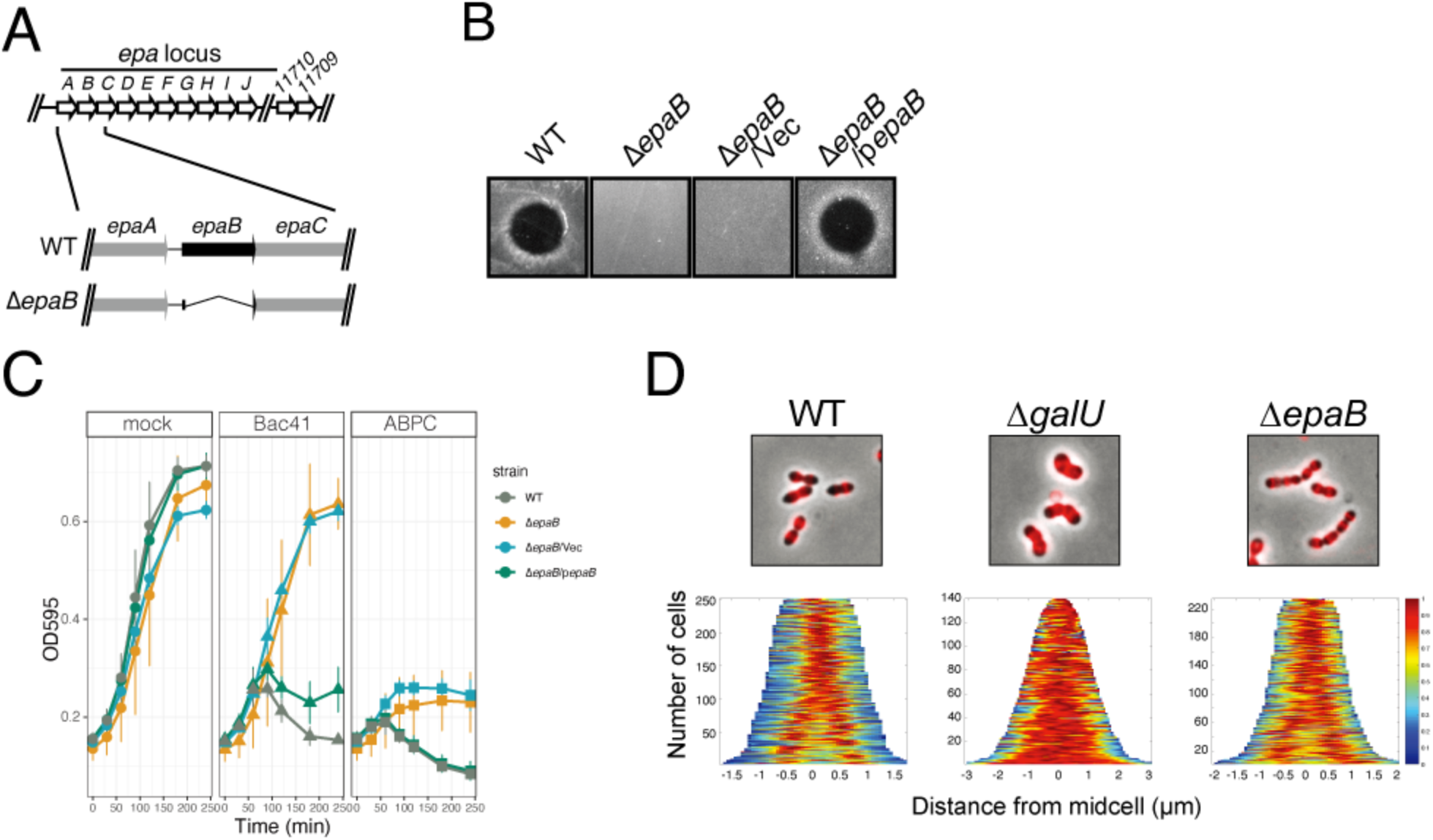
Effect of *epaB* deletion on the binding affinity of BacL_1_ and susceptibility to Bac41 activity. **(A)** Genomic organization of the *epa* locus in *E. faecalis* OG1 RF and construction of an epaB-deletion mutant. (B) A mixture of the recombinant BacL,-His and BacA-His proteins (25 ng each) was spotted onto THB soft agar (0.75 %) containing the indicator strain E. *faeca/is* wild type, *ΔepaB, ΔepaB/Vec* and *ΔepaB/pepaB* strains. The plate was incubated at 37 °C for 24 h, and the formation of halos was evaluated. (C) The overnight cultures of the *E. faecalis* strains OG1 RF (WT), *ΔepaB, ΔepaB/Vec* and *ΔepaB/pepaB* were inoculated into fresh THB broth at a 10-fold dilution in the presence or absence of ABPC (4 mg/L) or Bac41 (BacL_1_ and BacA, 2.5 μg/ml each), followed by incubation at 37 °C. The turbidity (OD_595_) was monitored during the incubation period. The data for each case are presented as the % of the initial turbidity. The data are presented as the mean ±8.D. (error bars) of three independent experiments.(D) Overnight cultures of the E. *faeca/is* wild-type, *Δga/U* and *ΔepaB* strains were diluted 5-fold with fresh THB broth and incubated with Hilyte Fluor 555 fluorescent dye-labeled (red) Bacl_1__SH3-His (5 μg/ml), followed by analysis via fluorescence microscopy.

To test whether BacL_1_ degradation susceptibility of cell wall lacking EPA, cell wall purified from the *ΔepaB* mutant was subjected to cell wall degradation assay (Supplementary figure S5). BacL_1_ was able to degrade cell wall of all tested strains including the *ΔepaB*, *ΔgalU* (*galU* in-frame deletion) and their derivatives with plasmids for complementation as well as wild-type strain. In addition, BacL_1_ could bind to cell surface of the *ΔepaB* mutant as wild type (Fig. S3B). In order to characterize localization of BacL_1_ binding on each strain in detail, we observed relative subcellular-localization of fluorescein-labeled SH3-repeat of BacL_1_ (BacL_1__SH3), which expresses strong and stable fluorescence signal compared to full-length BacL_1_ protein (20), on wild type, the *ΔgalU* and the *ΔepaB* mutants (Fig. 5D). On demograph based on microscope image, BacL_1__SH3 bound around division site of *ΔgalU* and *ΔepaB* eually to wild type as previously reported (20). It is noted that BacL_1__SH3 localization in the *ΔgalU* mutant was even stronger and more dispersed on the cell surface comapred to wild type. These observations are in line with our previously mentioned hypothesis that alternation of cell wall integrity does not affect BacL_1_ targeting for endopeptidase activity and does not underlie resistant phenotype of these mutant against Bac41-indused cell lysis.

### The *galU* inactivation and loss of EPA reduce susceptibility to beta-lactams

It has been suggested that EPA deficiency results in reduced cell surface integrity and consequently leads to increased susceptibility to antimicrobial agents or several environmental stress (31, 33, 34). To test this phenotype in the *galU*^−^ strain, we evaluated susceptibility to antibiotics in the OG1S derivatives (Table 2). Consistent with previous reports, bacteriostatic antibiotics (vancomycin and gentamycin) and detergent (sodium dodecyl sulfate (SDS)) were more effective against the *galU*^−^ and *galU*^−^/Vec strains than the wild-type and complemented strains. In contrast, the MICs of beta-lactams (ampicillin and penicillin G) for the *galU*^−^ strain were 4 mg/L and 2 mg/L, respectively, indicating reduced susceptibility in comparison to the parent strain. As already shown by Singh and Murray (35), the *ΔepaB* strain also displayed higher MICs and reduced susceptibility for beta-lactams than its parent (Table 2, Fig. 5C). The viability of the *galU*^−^ mutant after treatment with ampicillin (ABPC; 4 mg/L) for 4 h was increased approximately 10-fold in comparison to the wild-type strain (Fig. 6A). In microscopic experiments, the wild-type strain displayed morphological changes, with an elongated or rhomboid shape after 60 min of treatment with ABPC (4 mg/L), and most cells exhibited drastic cell disruption between 90 and 180 min after the treatment (Fig. 6B). On the other hand, the *galU*^−^ strain displayed a milder phenotype, with a swollen shape, showing a response distinct from that of the wild-type strain (Fig. 6B). In addition, only a few disrupted *galU*^−^ cells were observed, even 180 min after treatment with AMP. To distinguish between a bacteriolytic effect and a growth inhibition (bacteriostatic) effect of AMP, the turbidity of bacterial broth cultures was monitored in the presence and absence of ABPC (4 mg/L) (Fig. 6C). In the wild-type strain, the turbidity was increased after 60 min of incubation, in the presence of ABPC and in the mock control. However, after 90 min, a drastic reduction of turbidity was detected, and the turbidity was eventually lower than the initial turbidity. In contrast, after exposure to AMP, the *galU*^−^ strain continued to grow constantly without reduced turbidity, although its growth rate was slower than that of the untreated culture. These results were consistent with the microscopic observations, in that the *galU*^−^ mutant could be susceptible to the bacteriostatic effect of ABPC but not to the bacteriolytic effect. Furthermore, fluorescently detection assay for cell lysis by Ethidium homodimer (EthD) clearly showed that ABPC treatment increased dead cell population in wild type (Fig. 6D and E). On the other hand, the *ΔgalU* and *ΔepaB* mutants did not show significant increases of dead cell population compared to un-treatment control. Also, cell elongation effect of ABPC is relatively less in the *ΔgalU* and the *ΔepaB* mutants while ABPC treatment resulted in the significant elongation of cell length in wild type (Fig. 6F). Altogether, these data suggested that unlike other drugs and detergents, the bacteriolytic action of beta-lactams requires an intact cell surface composition constructed by the *galU* or *epa* genes.

**Figure 6.**
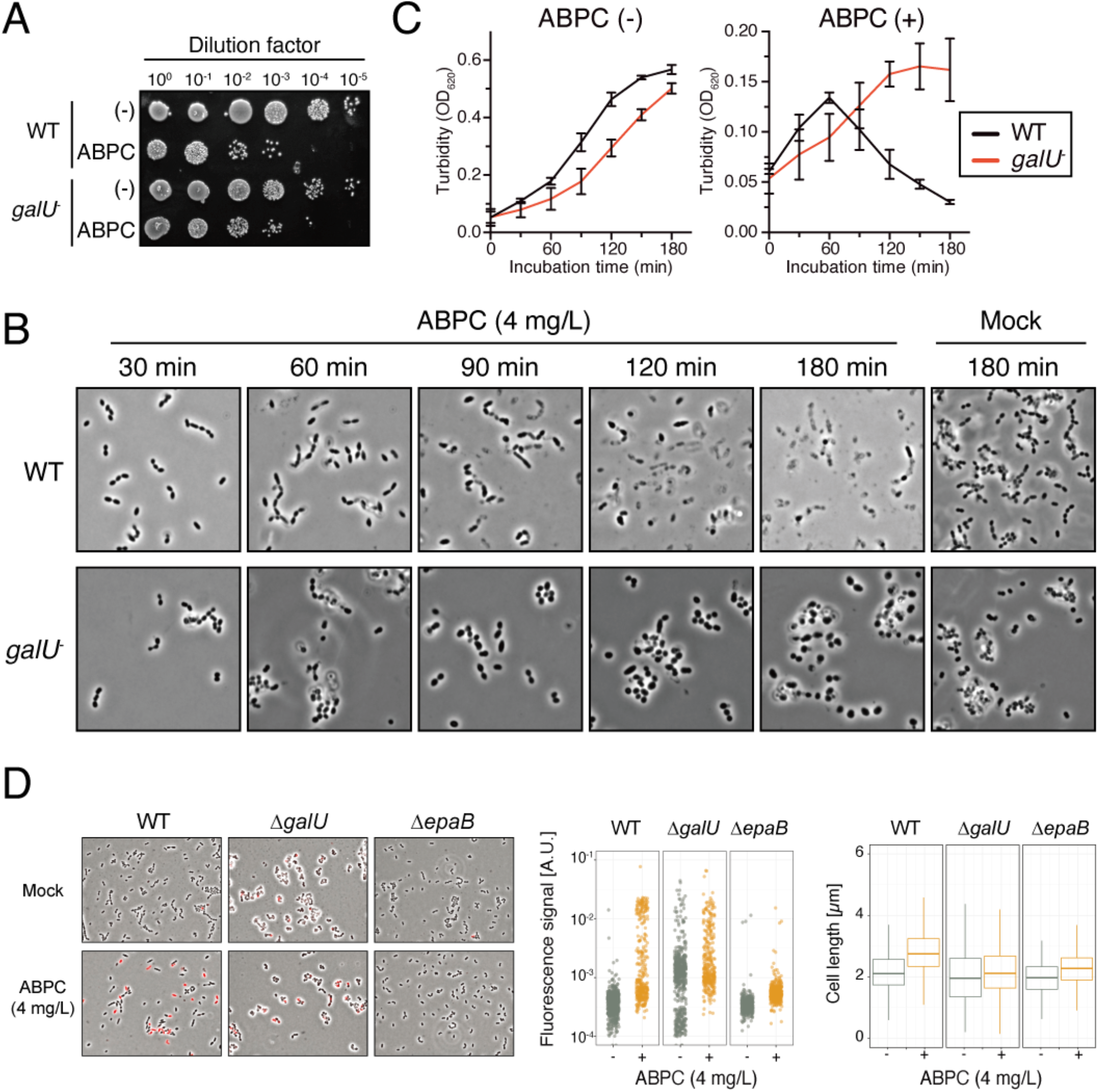
Different bacteriolysis phenotypes induced by ampicillin treatment (ABPC) in the *E. faecalis* wild-type strain or the *ga/U* mutant. (A) The *E. faecalis* strains OG1S (WT) and the *ga/U*^−^ mutant were treated with ABPC (4 mg/L) at 37 °C for 3 h. The bacterial suspensions were serially diluted 10-fold with fresh THB and then spotted onto a THB agar plate, followed by incubation overnight. Colony formation was evaluated as a measure of bacterial viability. (B) Confluent cultures of the *E. faecalis* OG1S (WT) and *ga/U*^−^ strains were diluted 10-fold with fresh THB containing ABPC(4 mg/L), followed by incubation at 37°C. The bacterial suspension was mounted onto a slide and analyzed via microscopy (phase contrast) at the indicated time points: 30, 60, 90, 120 and 180 min after treatment. The yellow arrowheads indicate the cell debris generated by ABPC-induced bacteriolysis. Cells incubated under identical conditions except for the absence of ABPC (Mock) are represented in the right panel as a reference. Scale bar, 20 *μm.* (C) The overnight cultures of the *E. faecalis* strains OG1S (WT) and *ga/U*^−^ were inoculated into fresh THB broth at a 10-fold dilution in the presence or absence of ABPC(4 mg/L), followed by incubation at 37 °C. The turbidity was monitored during the incubation period. The data for each case are presented as the % of the initial turbidity. The data are presented as the mean ±S.D. (error bars) of three independent experiments. (D) The overnight cultures of the *E. faecalis* strains OG1 RF (WT), *ΔgalU* and b. *epaB* were inoculated into fresh THB broth at a 10-fold dilution in the presence or absence of ABPC (4 mg/L), followed by incubation at 37°C for 1 h with EthD (2 mg/L). The bacterial suspension was mounted onto a slide and analyzed via fluorescence microscopy. Images are shown as merge of phase contrast and red fluorescence signal (EthD). (E and F) Quantification of population with red fluorescence intensity (E) and cell length (F) on each celll was generated images of based on panel D.

## Discussion

### The function of the *E. faecalis* GalU C-terminus

Here, we demonstrated that a spontaneous Bac41-resistant isolate carried a truncation mutation in the *galU* gene, with the deletion of 11 C-terminal residues (Fig. 1). Like related species, the GalU of *E. faecalis* contains a conserved nucleotide transferase domain at 5–257 a.a (Supplementary figure S6 and S7). However, the resistant mutant has a truncation deletion of 288–298 a.a., which does not correspond to the conserved domain. Despite this anomaly, this partial truncation leads to the inactivation of the protein’s UDP-glucose phosphorylase activity (Fig. 2 and Supplementary figure S1). Structural studies of GalU homologues from several bacterial species revealed that their C-terminal domains entangled each other to form homo-dimers (36, 37). Thus, it is possible that the deletion of just 11 amino acids from the C-terminal moiety might impair this dimerization, resulting in a conformational defect that affects GalU enzymatic activity.

### The role of *galU* in cell wall polysaccharide production in *E. faecalis* strains

The *gpsA*-*galU* locus is conserved among *Enterococcus* and *Streptococcus* species, although its chromosomal location and flanking genetic context are unrelated (Supplementary figure S7). This locus has been best studied in *Streptococcus pneumoniae*, which is a bacterial pathogen that causes pneumonia in humans (38). The surface capsular polysaccharide (CPS) of *S. pneumoniae* is a major virulence factor and the target for vaccination. More than 90 CPS types that have been defined in *S. pneumoniae* are synthesized from the corresponding specific *cps* locus, which is located between *pbp2X* and *pbp1A* (39). Although the *cps* genes are variable and specific for each of the 90 capsule types, *gpsA*-*galU* is highly conserved in a location that is distant from *cps* and is essential for the biosynthesis of every capsule type (40). Thus, *gpsA*-*galU* is suggested to play a critical role in the highly conserved universal pathway for cell surface-associated polysaccharides in various lactic acid bacterial species (41, 42). For *E. faecalis*, four serotypes (A, B, C and D) have been identified (43). These serotypes are defined by variants of two cell surface components: CPS and EPA (44). The CPS is serotype-specific and is represented only in serotypes C and D but not in serotypes A and B (28). Conversely, EPA is represented in every *E. faecalis* strain with any serotype. OG1 strains belong to the serotype B lineage that possesses only EPA. Meanwhile, the serotype C strains, such as FA2-2 and V583, produce both EPA and CPS (43, 44). Teng *et al*. identified the EPA-synthesizing *epa* gene cluster (locus tag 11715–11738), which is distant from the *gpsA*-*galU* locus (locus tag 11457 and 11458), on the OG1RF chromosome (29, 31). Recent study by Guerardel *et al*. defined structure of EPA and it consists of several sugars, including glucose, galactose, rhamnose, and ribitol (27). According to previous studies of capsular polysaccharide synthesis, those sugars are utilized to construct the cell wall glycopolymer through uridyl-nucleotide intermediates, such as UDP-glucose, which is produced by GalU. The data showing that the inactivation of the *galU* gene caused the abolishment of EPA (Fig. 2C) demonstrated the involvement of the *gpsA*-*galU* locus in the universal biosynthetic pathway for cell surface polysaccharide species in *E. faecalis*, which is similar to available reports for *S. pneumoniae*. The deletion of *galU* resulted in the complete loss of cell wall-associated polysaccharides in the serotype C *E. faecalis* FA2-2 (Supplementary figure S8), suggesting that GalU also plays a role in CPS production. This study is the first report to describe the function of *galU* in the biosynthesis of cell wall-associated polysaccharides in *E. faecalis*.

### Deduced model of *galU*-dependent susceptibility to lytic agents

A previous study revealed that the inactivation of *galU* decreases cellular robustness against several stresses, such as antibiotics and H_2_O_2_ (45). Here we confirmed that the *galU*^−^ strain displayed lower MICs for gentamicin, daptomycin and SDS in comparison to the parent strain (Table 2). However, we focused on the opposite phenotype that the inactivation of *galU* conferred resistance to the lytic bacteriocin Bac41 (BacL_1_ and BacA) and beta-lactams (Fig. 1, Fig. 6 and Table 2). In comparison with the wild-type strain, the *galU*^−^ strain produced a cell wall with slightly reduced susceptibility to BacL_1_ degradation (Fig. 4A). The contribution of this phenotype to resistance should be partial, and this process did not appear to be the fundamental mechanism by which the bacteriolytic activity of Bac41 is triggered by the undefined action of BacA following peptidoglycan degradation by BacL_1_ (20). UDP-glucose is essential for the biosynthesis of not only cell surface polysaccharides but also other cell wall-associated components, such as teichoic acid. The *ΔepaB* mutant that displayed a specific EPA defect conferred Bac41 resistance, as observed for the *galU*^−^ mutant (Fig. 5C). However, we cannot conclude that EPA contributed exclusively or directly to susceptibility to Bac41 because both mutants displayed the characteristic phenotype of abnormal cell morphology (Fig. 3 and Fig. 5) (30), suggesting that this drastic phenotype is caused by the loss of envelope integrity that results from the depletion of cell wall-associated polysaccharides (31). It is worth noting that the morphological defects observed in the *galU* and *epaB* mutants were not identical, which implies that other UDP-glucose-derived components might affect the phenotype of the *galU* mutant. In *Bacillus subtilis*, which is a model organism for Fermicutes, GtaB (a GalU homologue) is indispensable for cell morphology. The GtaB-deficient mutant exhibits a dislocation of FtsZ, which is a conserved protein that determines the contracting (separating) site during binary cell division, and the mutant consequently exhibits impaired cell division with an abnormal cell shape(46). As shown in Fig. 4, the *galU*^−^ mutant gave rise to a dispersed localization of BacL_1_, which binds specifically to the nascent cell wall and displays a limited localization at the mid-cell in the wild-type strain (20). Since the cell division machinery complex performs dual functions (synthesize and degrade peptidoglycan), the failure to properly control this machinery, especially the inhibition of penicillin binding proteins (PBPs), leads to lethal effects on bacterial cells. The potential intrinsic lethality of the cell division machinery has been suggested to underlie beta-lactam-induced bacteriolysis (47). Therefore, it is possible that the deactivated cell division machinery in the *galU*^−^ and Δ*epaB* mutants reduces the intrinsic lethality of the cell division machinery, resulting in resistance to Bac41 or beta-lactams. Collectively, this study provides insights into the intrinsic factors involved in the extrinsic bacteriolysis mechanism in *E. faecalis*, but further investigation is required to understand the precise mechanism that underlies these effects.

## Experimental procedures

### Bacterial strains, plasmids and antimicrobial agents

The bacterial strains and the plasmids used in this study are shown in Table 1. *Enterococcal* strains were routinely grown in Todd-Hewitt broth (THB) (Difco, Detroit, MI) at 37 °C (48), unless stated otherwise. *Escherichia coli* strains were grown in Luria-Bertani (LB) (Difco) medium at 37 °C. The antibiotic concentrations used to select *E. coli* were 100 mg/L of ampicillin (AMP), 30 mg/L of chloramphenicol and 500 mg/L of erythromycin. The concentrations used for the routine selection of *E. faecalis* harboring pMGS100 or pMSP3535 derivatives were 20 mg/L of chloramphenicol or 10 mg/L of erythromycin, respectively. Nisin was added at a concentration of 25 mg/L for the cultivation of the *E. faecalis* strain harboring pMSP3535. All antibiotics were obtained from Sigma Co. (St. Louis, MO).

**Table 1.**
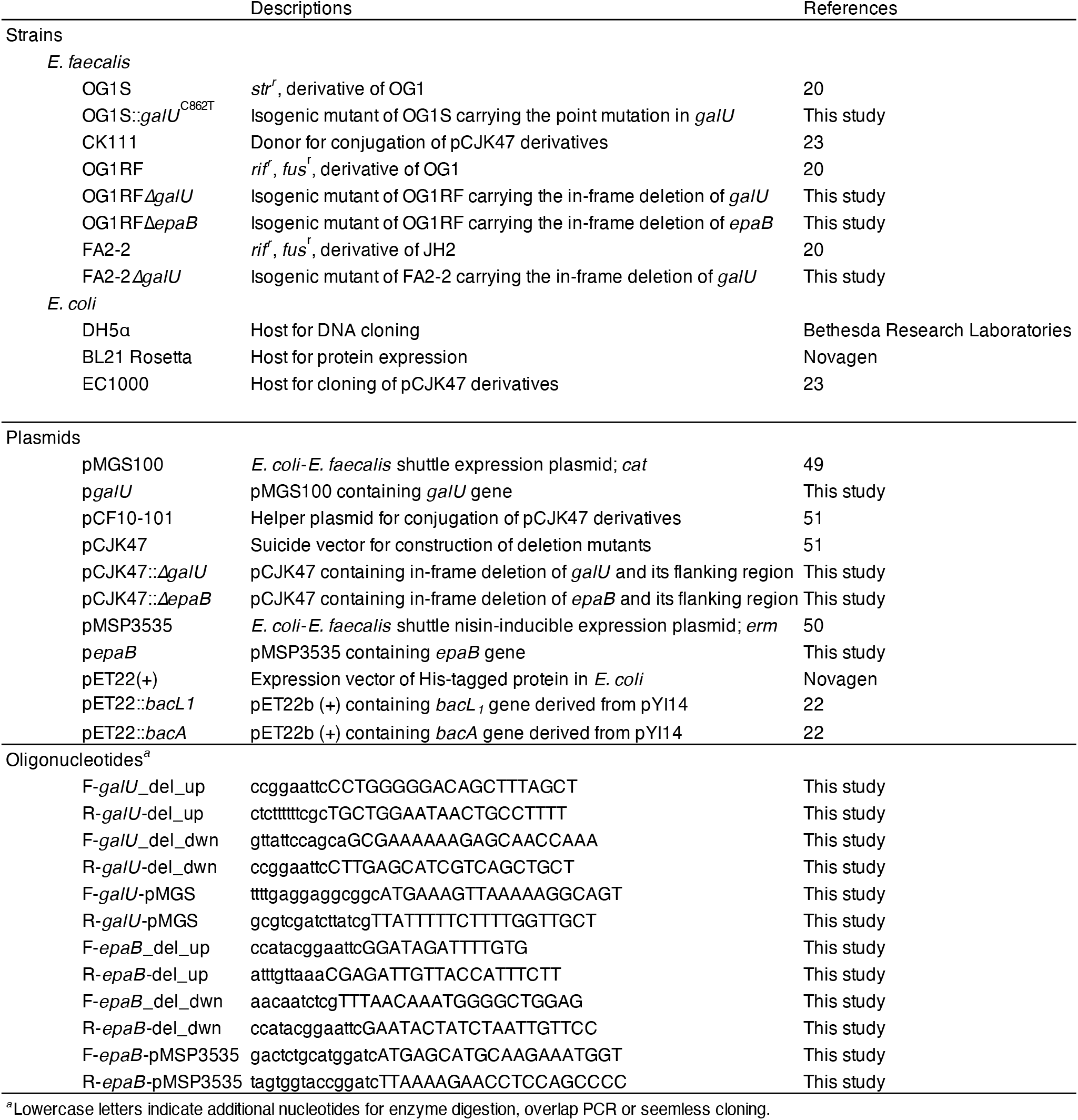
Bacterial strains and plasmids used in this study

**Table 2.**
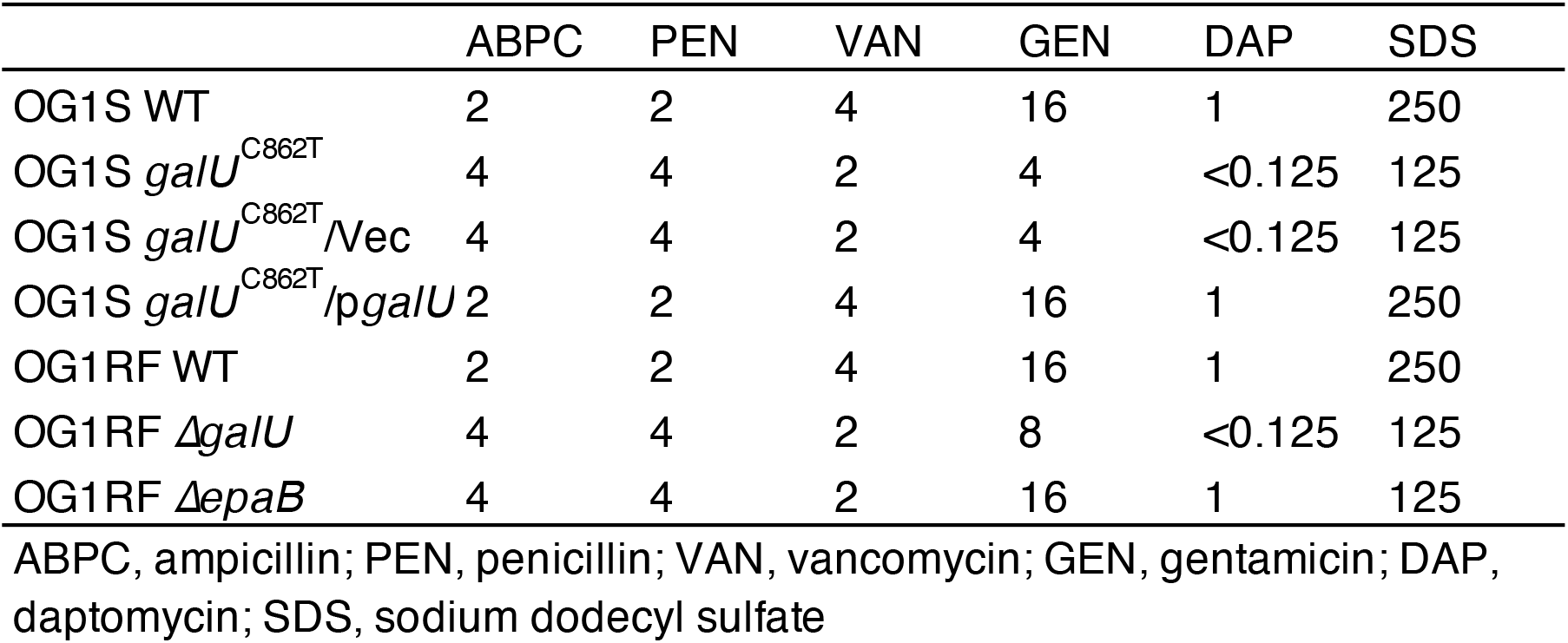
MICs of antimicrobial drugs for *E. faecalis* strains (mg/L)

### Soft-agar bacteriocin experiment

The soft-agar assay for bacteriocin activity was performed as described previously (7). Briefly, 1 μL of recombinant protein solution (25 ng/μl) was inoculated into THB soft-agar (0.75 %) containing the indicator strain and was then incubated at 37 °C for 24 h. The formation of an inhibitory zone was evaluated as sign of bacteriocinogenic activity of the test strain. The histidine-tagged recombinant proteins of BacL_1_ and BacA were prepared by the Ni-nitrilotriacetic acid (Ni-NTA) system (Invitorogen, Carlsbad, CA) as previously described (22).

### Next generation sequencing and variant analysis

Genomic DNA (gDNA) was extracted from *E. faecalis* OG1S and its isogenic spontaneous mutant using an Isoplant DNA isolation kit (Nippon gene, Tokyo, Japan) according to the manufacturer’s instructions, and submitted to Otogenetics Corporation (Norcross, GA USA) for exome capture and sequencing. Briefly, gDNA was subjected to agarose gel electrophoresis and OD ratio tests to confirm the purity and concentration of the DNA prior to Bioruptor (Diagenode, Inc., Denville, NJ USA) fragmentation. Fragmented gDNAs were tested for size distribution and concentration using an Agilent Bioanalyzer 2100 or Tapestation 2200 and Nanodrop. Illumina libraries were made from qualified fragmented gDNA using SPRIworks HT Reagent Kit (Beckman Coulter, Inc. Indianapolis, IN USA, catalog# B06938) or NEBNext reagents (New England Biolabs, Ipswich, MA USA, catalog# E6040) and the resulting libraries were then sequenced on an Illumina HiSeq2000/2500 which generated paired-end reads of 100 nucleotides (nt). Data was analyzed for data quality using FASTQC (Babraham Institute, Cambridge, UK).

### Construction of expression plasmids

To complement the *galU* defect in the *galU*^−^ strain, a *galU* expression plasmid designated as p*galU* was constructed as described below. A 927-bp fragment containing the full-length *galU* gene (897-bp) and an additional sequence at both ends (15-bp each) was amplified by PCR with the primers F-*galU*-pMGS and R-*galU*-pMGS, using *E. faecalis* OG1S genomic DNA as the template. Using In-fusion HD cloning (Clontech, Mountain View, CA), the resulting fragment was cloned into pMGS100 (49), which was linearized by digestion with *Nde*I and *Xho*I. The resulting plasmid was designated p*galU*. To complement the *epaB* defect, an *epaB* expression plasmid designated as p*epaB* was constructed as described below. An 819-bp fragment containing the full-length *epaB* gene (OG1RF_11737, 789-bp) and an additional sequence at both ends (15-bp each) was amplified by PCR with the primers F-*epaB*-pMSP3535 and R-*epaB*-pMSP3535, using *E. faecalis* OG1RF genomic DNA as the template. Using In-fusion HD cloning (Clontech, Mountain View, CA), the resulting fragment was cloned into pMSP3535 (50), which was linearized by digestion with *Bam*HI. The resulting plasmid was designated p*epaB*.

### Site-directed in-frame deletion of *galU*

Site-directed mutagenesis was carried out as described previously by Kristich *et al* (51). The 1-kbp flanking DNA fragments of the upstream or downstream regions of *galU* (OG1RF previously annotated *cap4C*) were amplified by PCR with the primers F-*galU*-del-up and R-*galU*-del-up or F-*galU*-del-dwn and R-*galU*-del-dwn, respectively, using *Enterococcus faecalis* OG1RF genomic DNA as the template. The resulting fragments were fused via overlapping PCR with the primers F-*galU*-del-up and R-*galU*-del-dwn, and the resulting 2-kb fragment was digested with *Eco*RI and cloned into pCJK47 at its *Eco*RI site to obtain pCJK47::*ΔgalU*. The pCJK47::*ΔgalU* plasmid was introduced into *E. faecalis* CK111 and was then transferred into *E. faecalis* OG1RF and FA2-2. The integrants of OG1RF and FA2-2 in which pCJK47::*ΔgalU* was integrated into the chromosome via primary crossing over were isolated through erythromycin selection. Cultivation without the drug allowed crossing over to facilitate the drop out of pCJK47 derivatives. The isolated candidates were screened by PCR with primers that were set outside of the region used for cloning into pCJK47. The site-directed deletion of the *epaB* gene was also carried out as described above, except pCJK47::*ΔepaB* was constructed using specific primer pairs (F-*epaB*-del-up and R-*epaB*-del-up or F-*epaB*-del-dwn and R-*epaB*-del-dwn) for the upstream region or the downstream region, respectively. Both the *ΔgalU* and *ΔepaB* mutants carry the initial 30 nt from the start codon and the terminal 30 nt, including the stop codon, without a frame-shift (N-terminal 10 a.a. and C-terminal 9 a.a.). The mutant genes encode the 19 a.a. fusion proteins with N-terminal 10 a.a. and C-terminal 9 a.a. regions.

### Scanning electron microscopy

A 200 μL overnight culture of bacteria was diluted 5-fold with 800 μL of fresh THB, and transferred onto a coverslip grass (Iwanami) in 24-well plate following incubation at 37 °C for 2 h (Iwanami). The coverslip was rinsed with 1 mL of 0.1M cacodylate buffer (pH 7.4). The bacteria on the coverslip were fixed with 1 mL of 1st fixation buffer [2% glutaraldehyde, 4% sucrose, 0.15% alcian blue, 0.1M cacodylate buffer (pH 7.4)] for 2 h at room temperature (RT). After primary fixation, the coverslip was washed three times with 0.1M cacodylate buffer (pH 7.4). An additional fixation was carried out with 1 mL of 2nd fixation buffer [0.5% OsO4, 0.1M cacodylate buffer (pH 7.4)] for 2 h at RT. The coverslip was washed with 0.1M cacodylate buffer (pH 7.4). The sample was dehydrated using an ascending ethanol series [50 % (1 min), 70% (2 min), 80% (3 min), and 100% (5 min ×2)] and was air-dried. Osmium coating was carried out at 5–6 mA, for 20 sec using an Osmium coater (Neoc-ST, Meiwafosis CO., LTD, Tokyo, Japan). The sample was observed in the Scanning electron microscopy (S-4100, Hitachi, Tokyo, Japan).

### Fluorescent microscopy

The red fluorescent dye-labeled recombinant protein of BacL_1_ was prepared with NH_2_-reactive HiLyte Fluor 555 (Dojindo, Kumamoto, Japan) as previously described (20). Bacteria diluted with fresh medium were mixed with fluorescent recombinant protein and incubated at 37 °C for 1 h. The bacteria were collected by centrifugation at 5,800 g for 3 min and then fixed with 4% paraformaldehyde at room temperature (RT) for 15 min. The bacteria were rinsed and resuspended with distilled water and mounted with Prolong gold antifade reagent with 4’, 6-di-amidino-2-phenylindole (DAPI; Invitrogen) on a glass slide. The sample was analyzed by fluorescence microscopy (Axiovert 200; Carl Zeiss, Oberkochen, Germany), and images were obtained with a DP71 camera (Olympus, Tokyo, Japan). For detection of dead cells, ethidium homodimer (EthD, Molecular probe) was added in bacterial culture at the final concentration of 2 ¼M and prepared for the microscopic analysis as described above. Raw image data were processed using FIJI(52). Cell segmentation and generation of fluorescence signal demograph were performed by Oufti(53). Statistical analysis and graphical representation on imaging data acquired by Oufti was performed by using R packages BactMAP(54) and ggplot2(55).

### Preparation and degradation analysis of the cell wall fraction

The bacterial culture was collected by centrifugation and rinsed with 1 M NaCl. The bacterial pellet was suspended in 4% SDS and heated at 95°C for 30 min. After rinsing with distilled water (DW) four times, unbroken cells were removed by centrifugation at 1,000 rpm for 1 min and the cell wall fraction in the supernatant was collected by centrifugation at 15,000 rpm for 10 min and was then treated with 0.5 mg/mL trypsin (0.1 M Tris-HCl [pH 6.8], 20 mM CaCl_2_) at 37 °C for 16 h. The sample was further washed with DW four times and was resuspended in 10 % trichloroacetic acid (TCA), followed by incubation at 4°C for 5 h, and then given four additional washes with DW. Finally, the cell wall fraction was resuspended in PBS. The cell wall degradation rate was quantified by measuring the optical density at 620 nm (OD620) using a microplate reader (Thermo).

### Preparation of cell surface polysaccharides

Cell surface polysaccharides were prepared as described by Hancock and Gilmore (44). An overnight bacterial culture was inoculated with as a 1:100 dilution into 25 ml of THB containing with 1 % glucose and grown at 37 °C for 5 h and then centrifuged to collect the cells (3,000 rpm, 10 min). The cells were washed with 2 ml of Tris-Sucrose solution [10 mM Tris (pH8.0), 25% sucrose]. The resulting cell pellets were resuspended in the Tris-Sucrose solution containing lysozyme (1 mg/ml) and mutanolysin (10 U/ml), and incubated at 37 °C for 16 h. The suspension was then centrifuged (8,000 rpm, 3 min). The supernatant was collected into a new tube and was treated with RNase I (100 μg/ml) and DNase I (10 U/ml) at 37 °C for 4h. Pronase E was added (20 μg/ml) and the sample further incubated at 37 °C for 16 h. 500 μl of chloroform was added to the sample, which was then centrifuged (12,000, 10 min). The aqueous phase (~300 μl) was transferred into a new tube and 920 μl of ethanol added (final concentration, 75%) to precipitate the contents at −80°C for 30 min. The precipitate was pelleted by centrifugation (15,000 rpm, 10 min), air dried, and resuspended with 100 μl of DW into one tube. Approximately 2.5 μl of the resulting sample was subjected to 10 % acrylamide gel electrophoresis buffered by TBE (10mM Tris, 10 mM borate, 2 mM EDTA). The separated carbohydrates were visualized with Stains-All (Sigma).

### Fermentation test

Fermentation of the respective sugar was examined as described before (40). Briefly, the *E. faecalis* strains were streaked on Heart infusion (HI, Difco) agar supplemented with 1% glucose or 1% galactose. Phenol red was added as a pH indicator at the final concentration of 25 ppm. After incubation overnight at 37 °C, fermentation was evaluated by the appearance of bacterial colonies and acidification of the agar medium.

### Antimicrobial susceptibility testing

The MICs of the antibiotics were determined by the agar dilution method according to CLSI recommendations [Clinical and Laboratory Standards Institute (http://clsi.org/)]. An overnight culture of each strain grown in Mueller-Hinton broth (Nissui, Tokyo, Japan) was diluted 100-fold with fresh broth. An inoculum of approximately 5×10^5^ cells was spotted onto a series of Mueller-Hinton agar (Eiken, Tokyo, Japan) plates containing a range of concentrations of the test drug. After incubation at 37 °C for 24 h, the number of colonies that had grown on the plates was determined.

## Acknowledgements

We thank Dr. Gary Dunny for providing the bacterial strains and the plasmids used to construct the mutants, Ms. Mari Arai and Ms. Satomi Tsuchihashi for providing technical assistance in this study, and Dr. Elizabeth Kamei for providing English proofreading of this manuscript.

---

**Supplementary figure S1.**
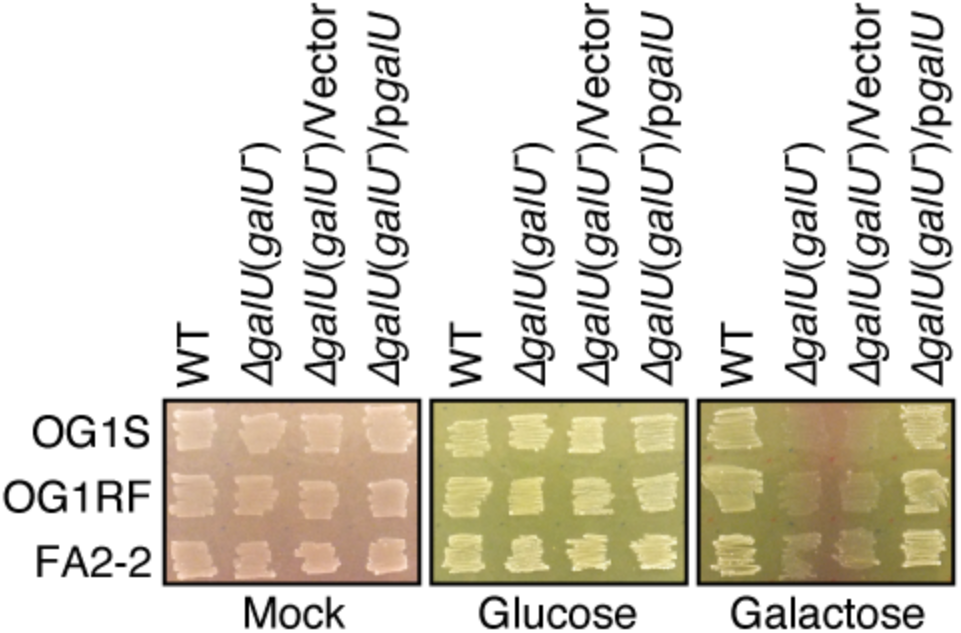
Effect of the *ga/U* inactivation on glucose or galactose fermentation. The indicated *E. faecalis* strains were grown in HI agar media that was supplemented with phenol red as a pH indicator and glucose or galactose as the fermentation source, respectively.

**Supplementary figure S2.**
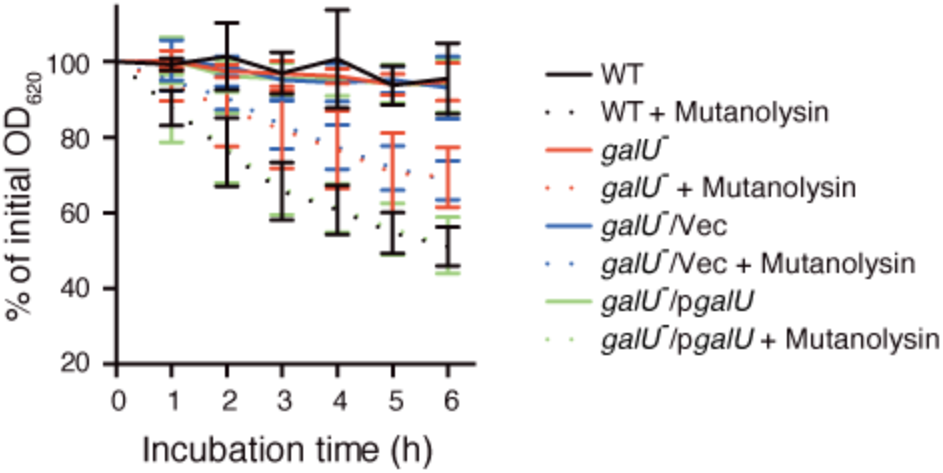
Susceptibility of cell wall derived from the *E. faeca/is* strains to Mutanolysin. Cell wall fractions prepared from *E. faecalis* wild type (black), *ga1U*^−^ (red), *ga/U*/*Vec* (blue) and *galU*^−^*lpga/U* (green) in exponential phase were diluted with PBS. Mutanolysin (1 μg/ml, dotted lines) was added to the cell wall suspension, and the mixture was incubated at 37^°^C. The turbidity at 620 nm was quantified at the indicated times during incubation. The present values are the percentages of the initial turbidity for the respective samples. The PBS treated sample (mock) is presented as a negative control (straight lines). The data are presented as the mean ±S.D. (error bars) of four independent experiments.

**Supplementary figure S3.**
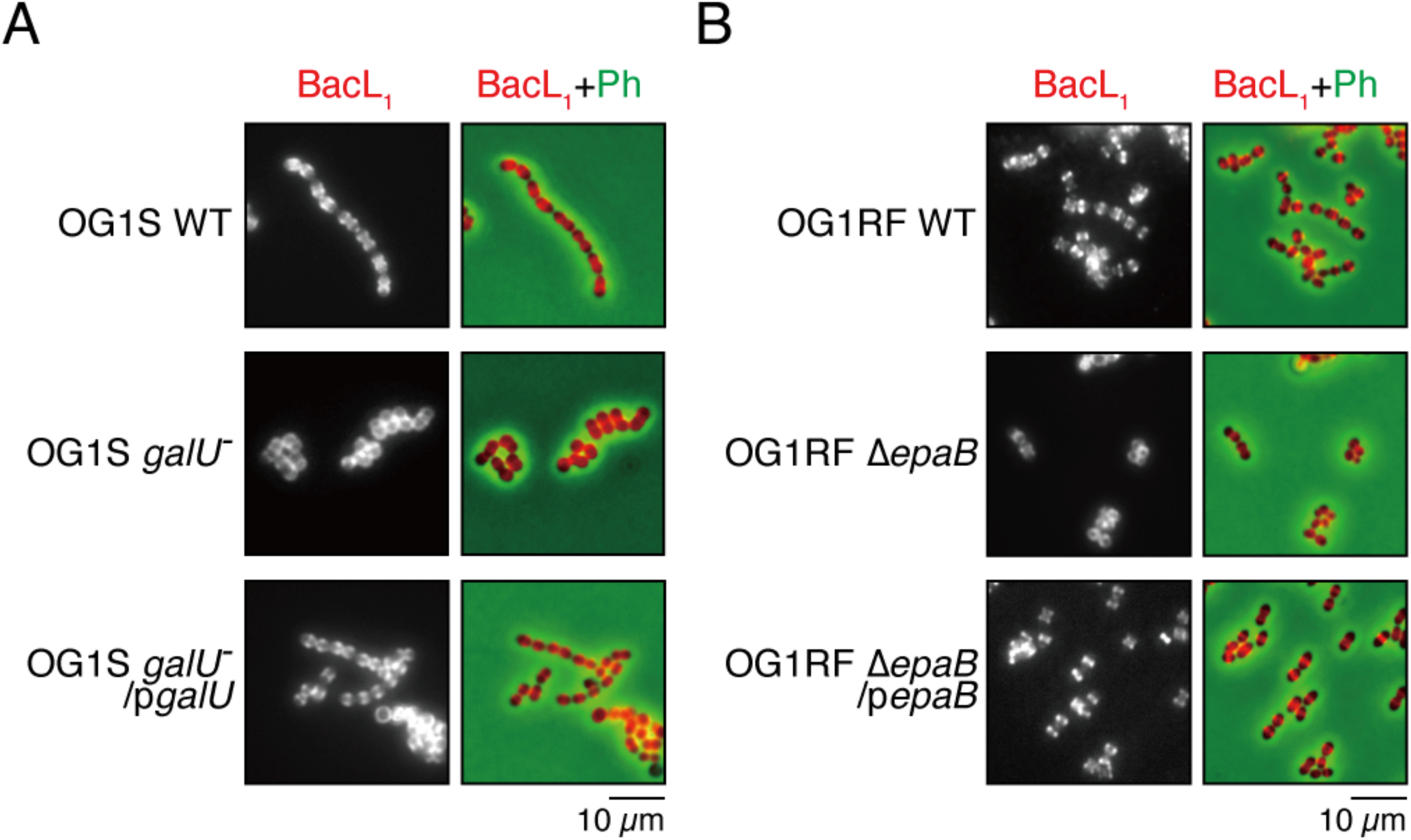
Overnight cultures of *E. faecalis* OG1 S wild type, *gaIU*^−^ and *gaIU*^−^*/pgaIU*^−^ strains (A) or *E. faeca/is* OG1 RF wild type, *ΔepaB* and *ΔepaB/pepaB* strains (8) were diluted 5-fold with fresh TH8 broth, and were incubated with the Hilyte Fluor 555 fluorescent dye-labeled (red) 8acL**1**(5 μg/ml), followed by analysis using fluorescence microscopy. Phase contrast (Ph) is pseudocolored (green) in merged images. These image in panel A or 8 are wide-field versions of the magnificated fluorescent images represented in Fig. 4B or Fig. 5D, respectively. Scale bar; 1 O *μm*.

**Supplementary figure S4.**
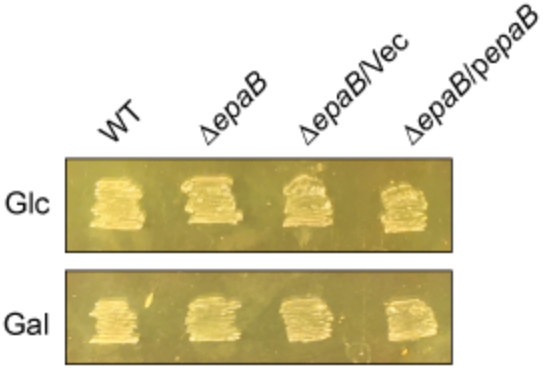
Effect of the *epaB* inactivation on glucose or galactose fermentation. The indicated E. *faecalis* strains were grown in HI agar media that was supplemented with phenol red as a pH indicator and glucose (Glc) or galactose (Gal) as the fermentation source, respectively.

**Supplementary figure S5.**
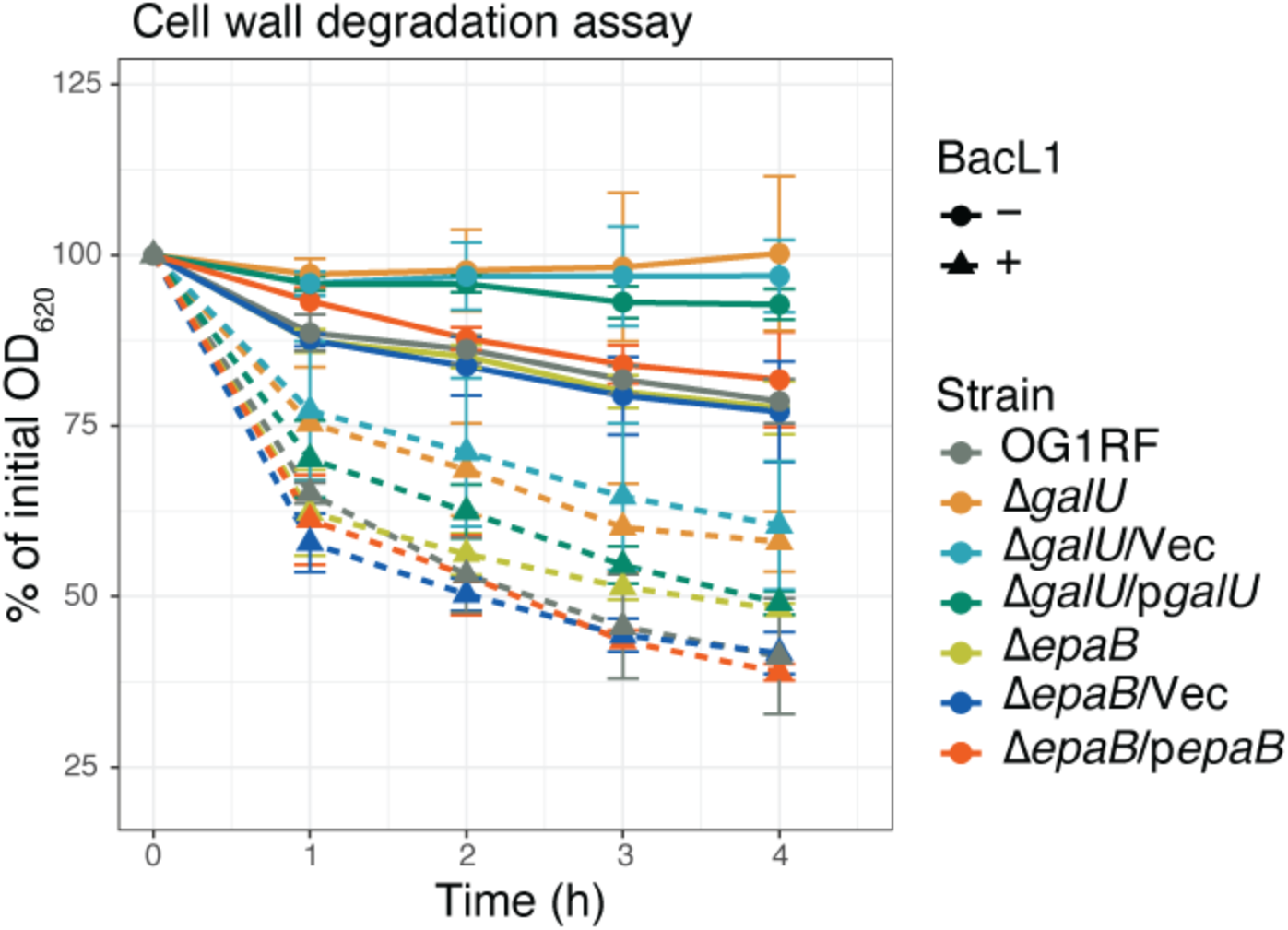
Susceptibility of cell wall derived from the *E. faecalis* strains to recombinant BacL,. Cell wall fractions prepared from *E. faecalis* wild type, *ΔgalU, ΔgalUNec, ΔepaB, ΔepaB/Vec* and *ΔepaB/pepaB* in exponential phase were diluted with PBS. BacL,-His (5 μg/ml) was added to the cell wall suspension, and the mixture was incubated at 37 C. The turbidity at 620 nm was quantified at the indicated times during incubation. The present values are the percentages of the initial turbidity for the respective samples. The PBS treated sample (mock) is presented as a negative control (straight lines). The data are presented as the mean ±S.D. (error bars) of four independent experiments.

**Figure S6.**
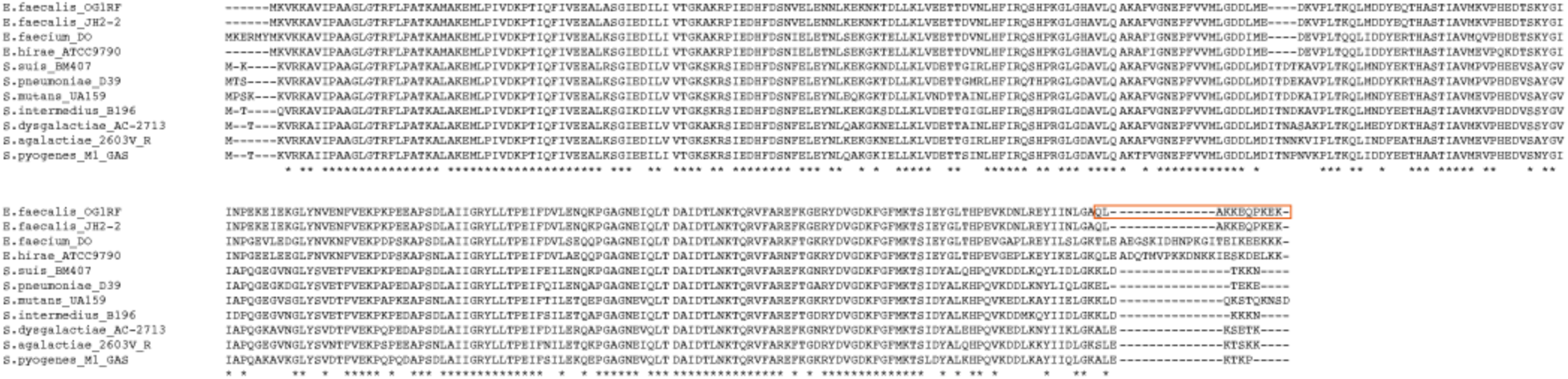
Alignment of the amino acid sequences of GalU homologues (GalU/HasC) from various Gram-positive bacterial species. The deleted region in the *ga/U-* strain used in this study is surrounded by a red frame. Source strains and GI numbers: *E. faecalis* OG1S/OG1R F (GI:327535310); *Enterococcus faecalis* J H2-2 (Gl:551092274); *Enterococcus faecium* DO (Gl:389869192); *Enterococcus hirae* ATCC9790 (Gl:498408730); *Streptococcus suis* BM407 (Gl:253756447); *Streptococcus pneumoniae* D39 (Gl:116076768); *Streptococcus mutans* UA159 (GI:24378821); *Streptococcus intermedius* B196 (GI:538452117); *Streptococcus dysgalactiae* AC-2713 (Gl:489151789); *Streptococcus agalactiae* 2603V R (Gl:22536589); and *Streptococcus pyogenes* M1 GAS (Gl:15674413).

**Supplementary figure S7.**
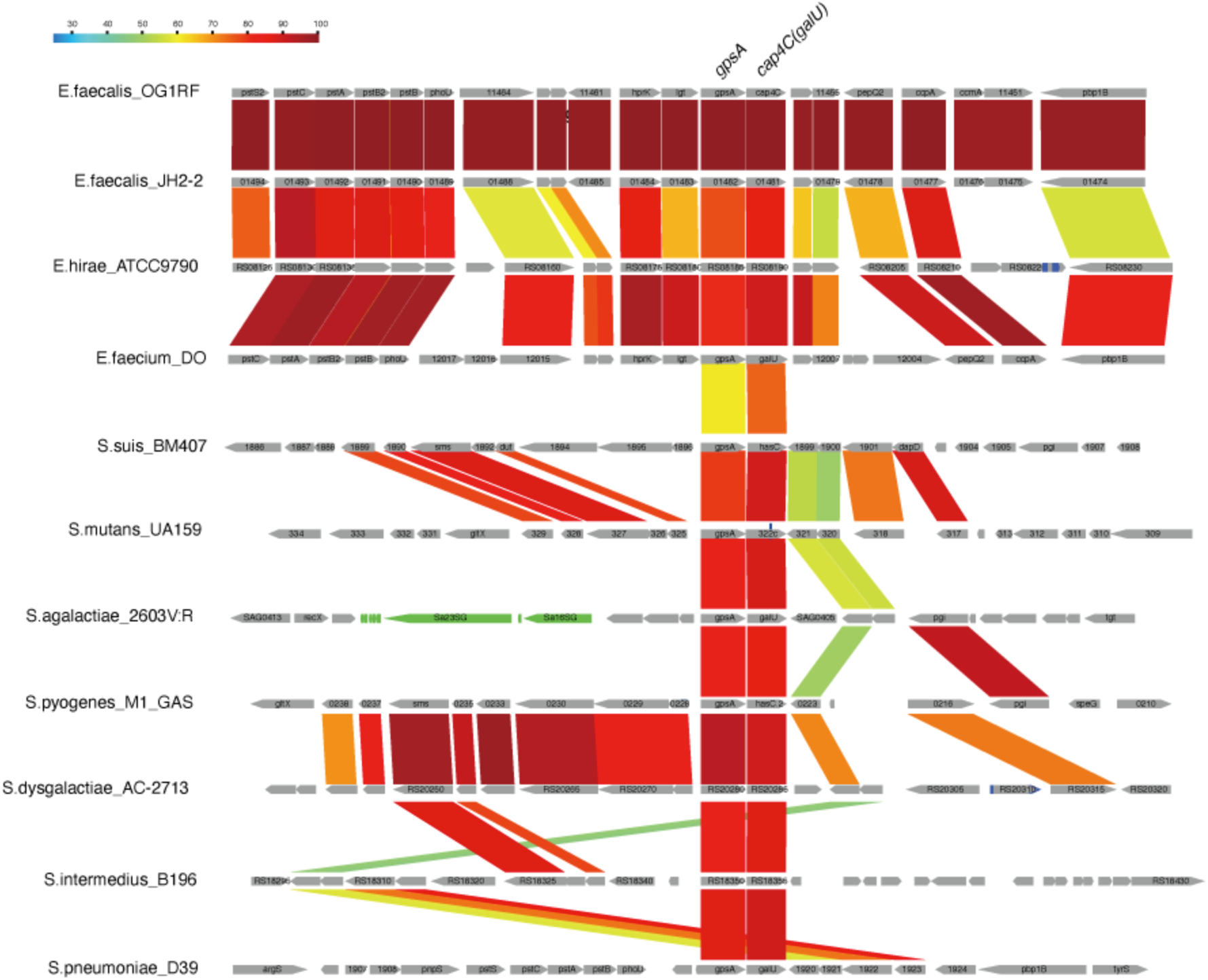
Genetic structure alignment for the flanking regions of *gpsA-ga/U* locus conserved among the Gram-positive bacterial species. Color scale represents similarity (%) of CDS based on the blastp analysis. The sequence data obtained from NCBI database (https://www.ncbi.nlm.nih.gov/). This scheme was generated using the GenomeMatcher software (http://www.ige.tohoku.ac.jp/oho/gmProject/gmhome.html) (1).

**Supplementary figure S8.**
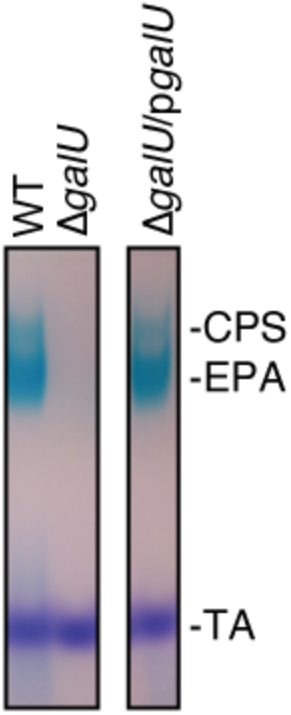
Effect of *ga/U* deletion on the cell wall-asociated polysaccharide production in ***E.** faeca/is* FA2-2. The ***E.** faecalis* FA 2-2 wild type, *b.ga/U*, and *b.ga/U/pga/U* strains were grown in THB broth supplemented with glucose, and the cell described in the Materials and Methods section. The resulting polysaccharides were separated via 10 % acrylamide gel electrophoresis, followed by staining with the Stains-All reagent. CPS, Capsule polysaccharide; EPA, Enterococcal polysaccharide antigen; TA, Teichoic acid.

